# A potent neutralizing epitope of limited variability in the head domain of haemagglutinin as a novel influenza vaccine target

**DOI:** 10.1101/212845

**Authors:** Craig P Thompson, José Lourenço, Adam A Walters, Uri Obolski, Matthew Edmans, Duncan S Palmer, Kreepa Kooblall, George W Carnell, Daniel O’ Connor, Thomas A Bowden, Oliver G Pybus, Andrew J Pollard, Nigel J Temperton, Teresa Lambe, Sarah C Gilbert, Sunetra Gupta

**Affiliations:** Department of Zoology, University of Oxford, UK; The Jenner Institute Laboratories, University of Oxford, UK; Oxford Centre for Diabetes, Endocrinology and Metabolism, University of Oxford, UK; Medway School of Pharmacy, University of Kent, UK; Oxford Vaccine Group, Department of Paediatrics, University of Oxford, and the NIHR Oxford Biomedical Research Centre, Oxford, UK; Division of Structural Biology, Wellcome Centre for Human Genetics, University of Oxford, UK.

## Abstract

Antigenic targets of influenza vaccination are currently seen to be polarised between (i) highly immunogenic (and protective) epitopes of high variability, and (ii) conserved epitopes of low immunogenicity. This requires vaccines directed against the variable sites to be continuously updated, with the only other alternative being seen as the artificial boosting of immunity to invariant epitopes of low natural efficacy. However, theoretical models suggest that the antigenic evolution of influenza is best explained by postulating the existence of highly immunogenic epitopes of limited variability. Here we report the identification of such an epitope of limited variability in the head domain of the H1 haemagglutinin protein. We show that the epitope mediates immunity to historical influenza strains not previously seen by a cohort of young children. Furthermore, vaccinating mice with these epitope conformations can induce immunity to all the human H1N1 influenza strains that have circulated since 1918. The identification of epitopes of limited variability offers a mechanism by which a universal influenza vaccine can be created; these vaccines would also have the potential to protect against newly emerging influenza strains.

## Introduction

Seasonal influenza is estimated to cause between 1-4 million cases of severe illness and 200,000 to 500,000 deaths per year^1^. The best way to protect against influenza infection is through vaccination. Currently, a trivalent (TIV) or quadravalent influenza (QIV) vaccine is given each year, targeting the circulating H1N1 and H3N2 influenza A strains and one or two lineages of the circulating influenza B strains. However, the vaccine has to be formulated at least 6 months prior to the influenza season and so the strains that are subsequently prevalent in the actual flu season do not always match the strains used in the vaccine^2^.

The antigenic evolution of influenza is known to occur through mutations in surface glycoproteins, principally haemagglutinin (HA), allowing strains to escape pre-existing host immunity^3-5^. Epitopes within HA are commonly assumed to be either highly variable due to strong immune selection (and typically located in the head domain of HA) or conserved due to the absence of immune selection (for example, in the stalk of HA)^6^. Together, these form the backbone of the theory of ‘antigenic drift’ whereby the virus population slowly and incrementally acquires mutations in protective highly variable epitopes. However, the antigenic drift model can only explain the epidemiology and limited genetic diversity observed among influenza virus populations when very specific constraints are placed on the mode and tempo of mutation or by invoking short-term strain-transcending immunity^7,8^. An alternative model known as ‘antigenic thrift’ successfully models the epidemiology and genetic diversity of influenza by assuming that immune responses against epitopes of limited variability drive the antigenic evolution of influenza^9-11^. Within this framework, new strains may be generated constantly through mutation but most of these cannot expand in the host population due to pre-existing immune responses against epitopes of limited variability. This creates the conditions for the sequential appearance of antigenically distinct strains and provides a solution to the long-standing conundrum of why the virus population exhibits such limited antigenic and genetic diversity within an influenza epidemic. An important translational corollary of this model is that a ‘universal’ influenza vaccine may be constructed by targeting such protective epitopes of limited variability.

We show that studies of sera from young children taken in 2006/7 using neutralisation assays and ELISAs reveals a periodic pattern of cross-reactivity to historical isolates consistent with the recycling of epitopes of limited variablity. We idenfity one epitope of limited variability responsible for this pattern through a structural bioinformatics analysis. We demonstrate that mutagenesis of the epitope removes the cross-reactivity to historical strains and vaccination of mice with the 2006 conformation of the epitope is able to reproduce the cross-reactivity pattern identifed in the serology studies. We further show that vaccination of other epitope conformations induces similar but asynchronous cross-reactivity to historical strains. Finally, we demonstrate that vaccination with the 2006 and 1977 epitope conformations is able to protect mice from challenge with a H1N1 influenza strain that last circulated in 1934. By establishing that the antigenic space within which influenza evolves is much smaller than previously thought, we show that there are epitopes in the major influenza antigen, HA, which if vaccinated against would allow us to avoid the requirement for yearly influenza vaccination, necessitated by the current TIV and QIV vaccines.

## Results

### Recognition of historical isolates follows a pattern consistent with recycling of epitopes

We tested the prediction that HA epitopes of limited variability exist by performing microneutralisation assays using pseudotyped lentiviruses displaying the H1 HA proteins from a panel of historical influenza isolates (hereafter described as pMN assays^12,13^); with sera obtained in 2006/2007 in the UK from 88 children born between March 1994 and May 2000 (Fig 1A). All individuals possessed neutralising antibodies to the A/Solomon Islands/3/2006 strain and 98% of individuals possessed neutralising antibodies to A/New Caledonia/20/1999, including those children born in 2000 who are unlikely to have been exposed to the strain. 99% of individuals also possessed neutralising antibodies to the A/USSR/90/1977 strain, while 30% of individuals also possessed neutralising antibodies to the A/WSN/1933 strain. By contrast, only 3%, 9.1% and 3.4% of individuals possessed antibodies to the A/California/4/2009, A/PR/8/1934 and A/South Carolina/1/1918 strains respectively. ELISA analysis using the HA1 domain of the same seven strains as an antigen was consistent with the pMN data and also identified broadly cross-reactive non-neutralising antibodies that bind the HA HA1 region of various H1 influenza strains (Fig S1A). These results are in agreement with number of recent studies suggesting antibody responses show some degree of periodic cross-reactivity to historical strains ^14-22^ counter to the view of ‘antigenic drift’ within which antigenic distance accumulates linearly with time.

**Figure 1:**
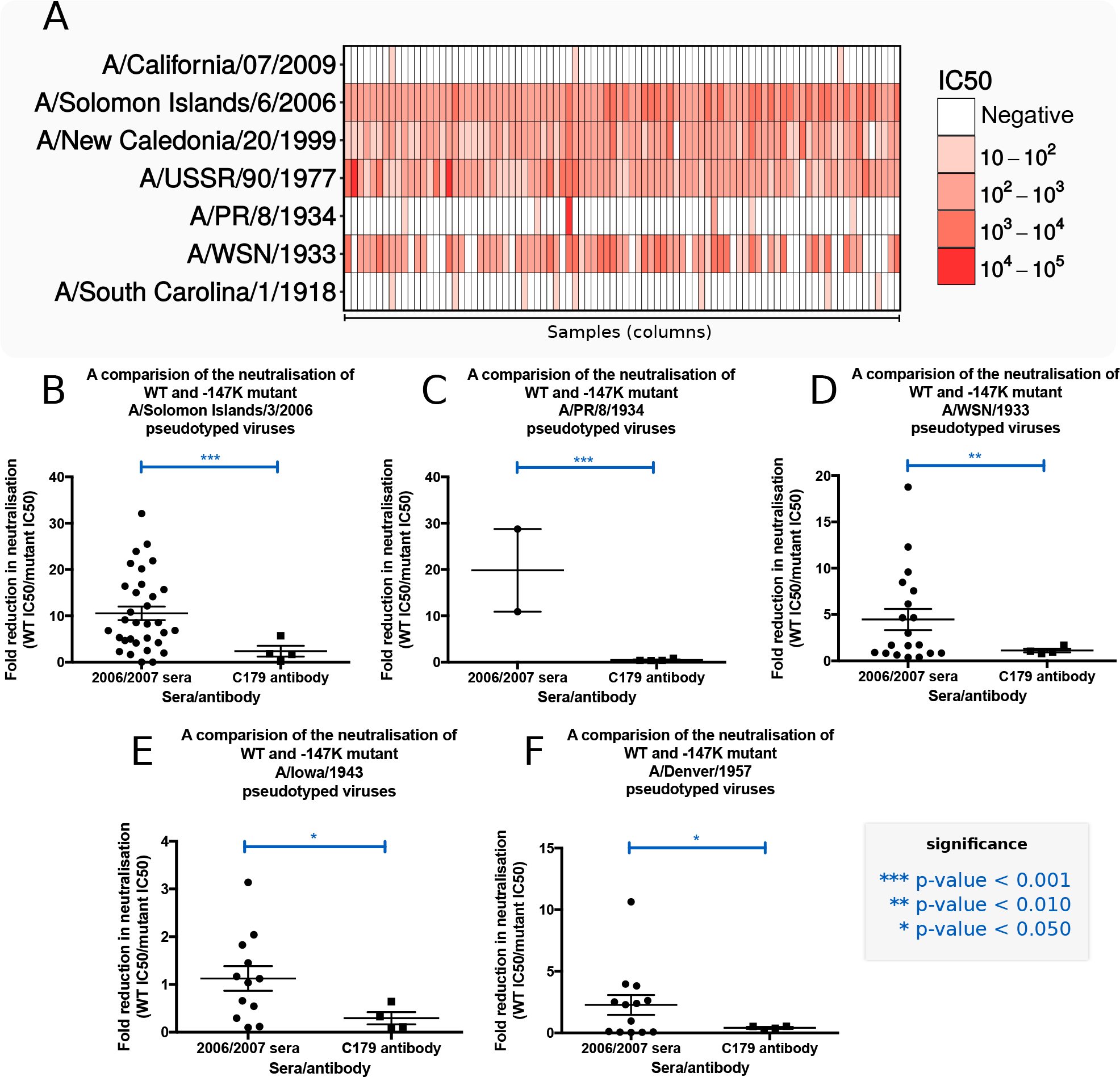
Pseudotype microneutralisation data revealing a cyclic pattern of epitope recognition and the involvement of position 147 in the production of cross-protective antibodies in sera taken from children aged 6 to 12 in 2006/2007. (**A**) Serum samples from children aged between 6 to 12 years in 2006/2007 n = 88 were tested for their ability to neutralise a panel of pseudotyped lentiviruses representing a range of historical isolates. (**B-F**) A lysine residue was inserted at position 147 linear numbering, where Met = 1 through site-directed mutagenesis SDM in the HAs of pseudotyped lentiviruses A/WSN/1933, A/PR/8/1934 and A/Solomon Islands/3/2006 included in panel A as well as A/Iowa/1943 and A/Denver/1957. The ratio of WT IC50 to mutant IC50 was then assessed to determine if there was reduction in neutralisation. A stalk targeting antibody, C179, was used as a control.

We noted that the A/Solomon Islands/3/2006, A/New Caledonia/20/1999, A/PR/8/34 and A/WSN/33 HAs all contained a deletion at position 147 (linear numbering, where Met = 1, H3 numbering: 133, WHO numbering: 130) and exhibited some degree of reactivity to the 2006/2007 sera. This position otherwise typically contains a positively charged arginine, in the case of the A/USSR/90/1977 HA, or lysine, as is the case for the A/California/4/2009 and A/Brevig Mission/1/1918 HAs. To determine whether the cross-reactivity observed between these strains could be attributed to the deletion, a lysine was inserted at position 147 (-147K) in the A/Solomon Islands/6/2006, A/PR/8/1934 and A/WSN/1933 HAs (Fig 1B-D). A statistically significant loss of neutralisation for the −147K A/Solomon Islands/3/2006, and A/PR/8/1934 and A/WSN/1933 mutant pseudotyped lentiviruses was observed (p-value= 0.0005, Fig 1B, p-value=0.0004, Fig 1C, and p-value=0.0056, Fig 1D). In the case of the A/PR/8/1934 −147K mutant, there was a total loss of neutralisation in four samples and a reduction in two samples indicating that the bulk of cross-reactivity between the A/Solomon Islands/3/2006 and the A/PR/8/1934 strains is mediated through an epitope located in the vicinity of the deletion (p-value= 0.0004, Fig. 1C). Therefore, it seems that the absence of a positively charged lysine at position 147 mediates much of the observed cross-reactivity to historical strains induced by the 2006/2007 cohort sera in Figure 1A.

Analysis of historical strain data shows that the deletion, the only one to occur in the H1 HA, appears periodically over the course to the antigenic evolution of H1N1 subtype of influenza, occurring in 1933, 1934, 1943, 1957 and between 1995 and 2008. To ascertain whether the absence of an amino acid at position 147 would also mediate cross-reactivity with other historical strains possessing the deletion, A/WSN/1933 neutralisation positive samples were run against the WT and −147K mutant A/Iowa/1943 and A/Denver/1957 pseudotyped lentiviruses (Fig 1E-F). A statistically significant reduction in neutralisation was observed for the −147K A/Iowa/1943 and A/Denver/1957 HA mutants (p-value= 0.012, Fig 1E, p-value= 0.011, Fig 1F). Furthermore, three samples, which neutralised the WT A/Denver/1957 HA, failed to neutralise the −147K mutant entirely. These results imply that at least part of the cross-reactive neutralising immune response within this cohort is mediated through the recognition of an epitope that contains a deletion at position 147. Moreover, the existence of a lysine at position 147 may contribute to the overall lack of neutralisation of A/California/4/2009 and A/South Carolina/1/1918.

### Identification of an epitope of limited variability

Several previous studies have highlighted the importance of position 147. Although not included within any of the canonical antigenic sites defined by Caton et al, 1982 (being absent in the A/PR/8/1934 Mt. Sinai strain), position 147 has recently been assigned to a new antigenic site denoted ‘Pa’ in Matsuzaki et al, 2014, where it was shown to be responsible for several A/Narita/1/2009 escape mutants^3,23^. Position 147 is also important for the binding of several known broadly neutralising antibodies: for example, the 5j8 antibody requires a lysine to be present at position 147, whilst the CH65 antibody cannot bind if a lysine is present at position 147^24,25^. Furthermore, Li et al, 2013 demonstrated that certain demographics, such as individuals born between 1983 and 1996, possess antibodies that bind to an epitope containing a lysine residue at position 147^18^.

We next employed a structural bioinformatic approach to identify an epitope of limited variability that contained position 147. *In silico* analysis was used to determine how the accessibility and binding site area contributed to the variability of hypothetical antibody binding sites (Fig 2A) on the surface of the A/Brevig Mission/1/1918, A/PR/8/1934, A/California/04/2009, A/Washington/5/2011 H1 HA crystal structures^26-30^. The antibody binding site (ABS) of lowest variability containing position 147 was consistently represented by the site shown in Figure 2B^31^, and could be shown to locate to an exposed loop in the head domain of the H1 HA, not covered by N-linked glycosylation (Fig 2B&C) and encompassing additional residues in the Ca_2_ antigenic site.

**Figure 2:**
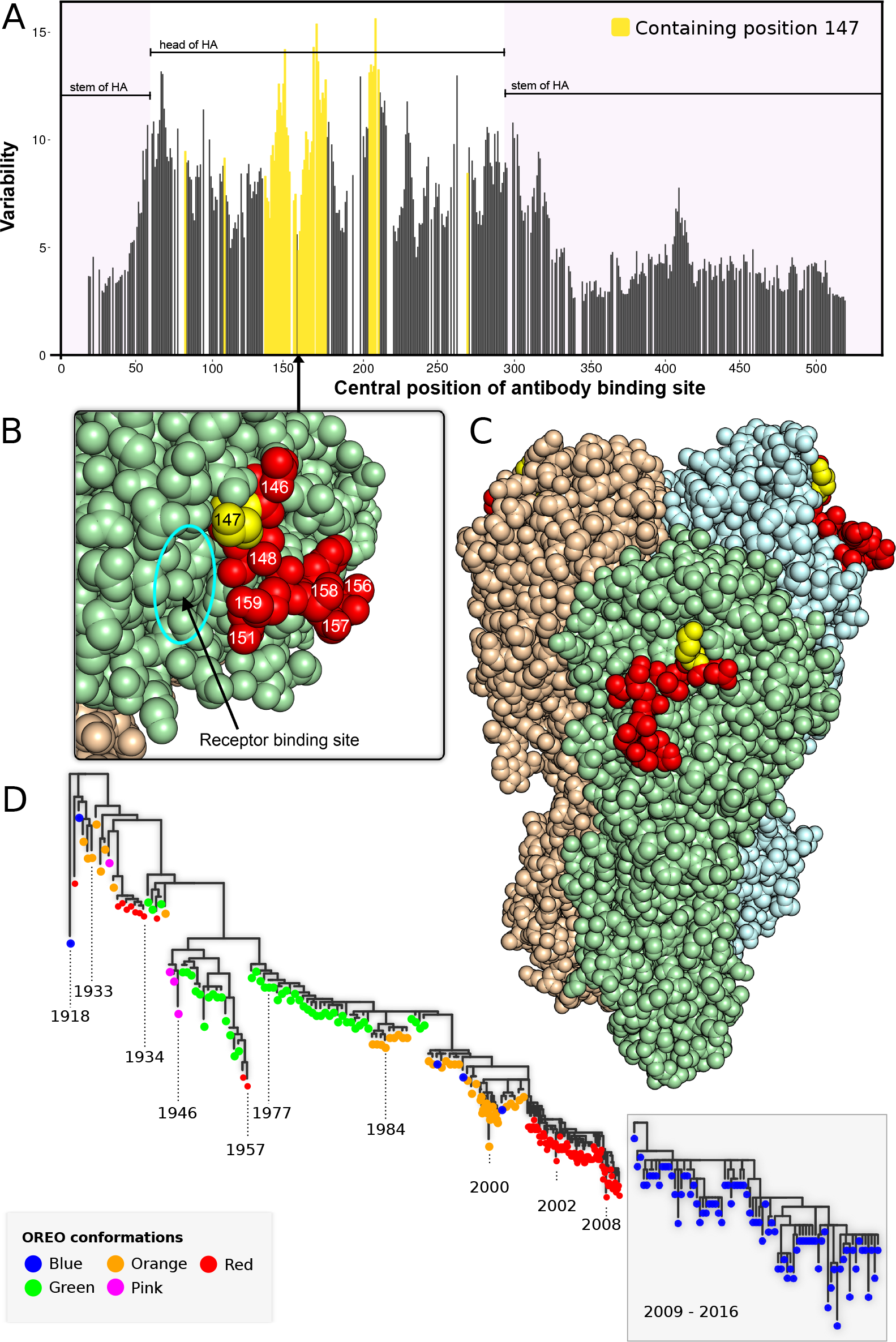
Identification of a site of limited variability in the head domain of the H1 HA through structural bioinformatic analysis. (**A**) Variability of antibody bindings sites ABS on the crystal structure of A/California/04/2009 HA; those containing position 147 are shown in yellow. (**B, C**) Location of ABS of lowest variability containing position 147 with position 147 shown in yellow and the rest of the site coloured in red. (**D**) Phylogenetic trees of pre-pandemic and post-pandemic highlighted rectangle H1N1 with tips coloured according to the conformation of the epitope of limited variability (hereafter called ‘OREO’).

Analysis of this site (hereafter called ‘OREO’) suggested that various conformational epitope variants could be defined on the basis of variation and structural proximity of positions 147, 158 and 159. Combining these analyses with the site directed mutagenesis SDM results, we arrived at a maximum of five epitope conformations of the epitope (Figs 3 & S3), which arise and disappear in a cyclical manner during the known evolutionary history of the pre-pandemic and post-pandemic H1N1 lineages (Fig 2D). This analysis demonstrates that there are numerous sites of limited variability in the head domain of the H1 HA, in addition to a range of highly variable sites (Fig 2A); the antigenic trajectory of the latter has been tracked in detail by several previous studies^32,33^.

**Figure 3:**
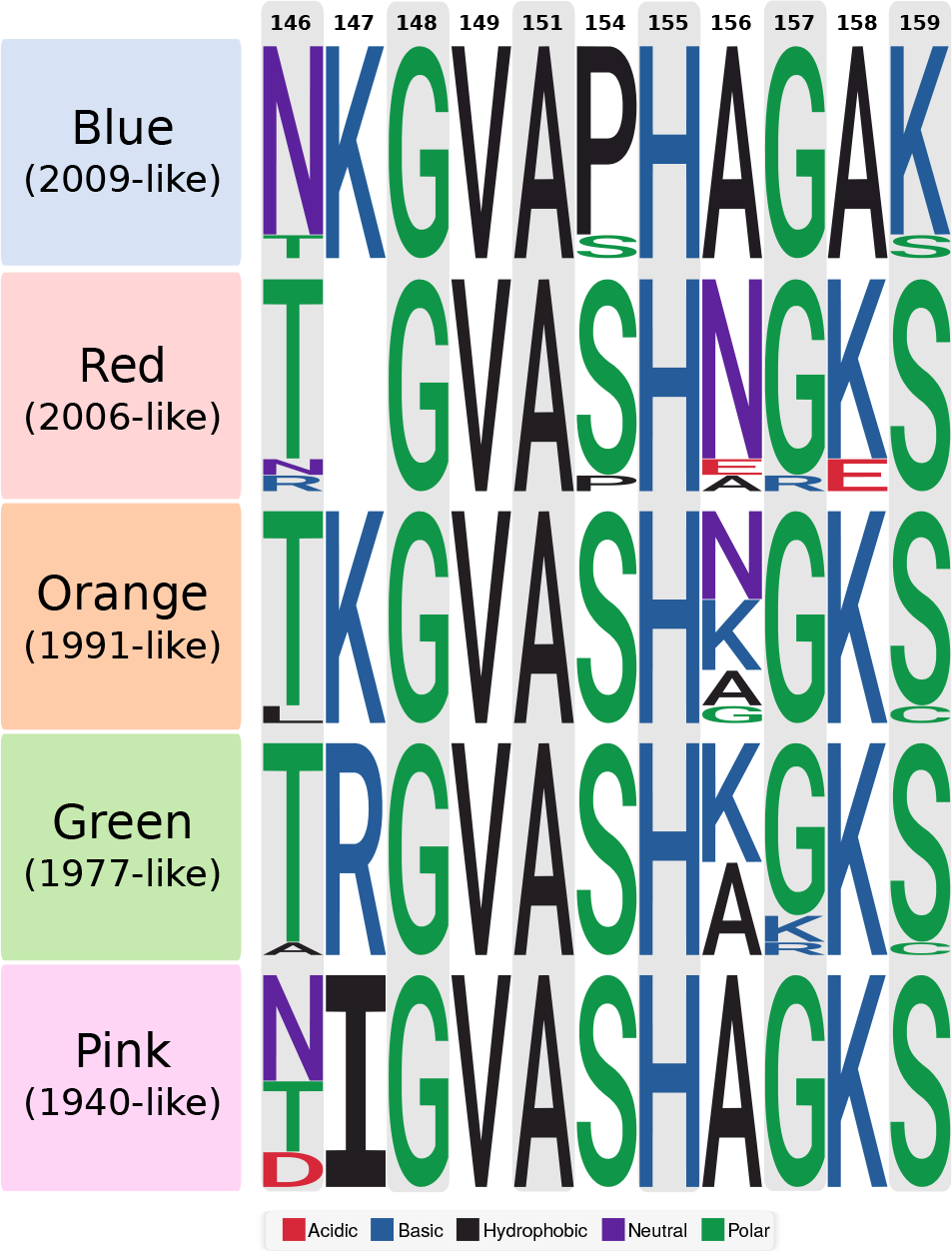
Allelic classes of the OREO epitope. Sequence Logo diagrams showing the relative frequency of different amino acids for each OREO conformation based on yearly consensus sequences. The various epitopes can be defined by the amino acids at positions 147, 158 and 159. These three positions have been used to define the conformations of OREO in the phylogenetic tree in Figure 2D.

### Vaccination with an epitope of limited variability induces cyclical cross-reactivity

We next substituted the five proposed conformations of OREO (Figs 3 & Table S1) into H5, H6 and H11 HAs, which have not circulated in the human population, to maintain the conformations of the epitope and allow the immune response to be focused on the epitope via a prime-boost-boost vaccination regimen (Figs 4A). Consequently, mice were vaccinated using a DNA-DNA-pseudotyped lentivirus regimen alternating between different HA scaffolds (Fig 4A). Analysis of sera obtained from the final bleed at 21 weeks prior to influenza challenge demonstrated that vaccinating with these epitopes produces antibodies that are cross-reactive to a number of historical strains. Notably, the 2006-like epitope conformation (red) produces cross-reactive antibodies that mirror the neutralisation profile of sera taken in 2006/2007 from young children aged 6 to 12 (Figs 1A&S1): both datasets show neutralisation of pseudotyped lentiviruses displaying HAs from A/Solomon Islands/3/2006, A/USSR/90/1977, A/PR/8/1934 and A/WSN/1933 via the OREO epitope but not A/California/4/2009 or A/South Carolina/1/1918 (Fig 4B-G). Intriguingly the 1977-like conformation (green), containing an arginine at position 147, also display similar cross-reactivity to that of the 2006-like epitope (red), containing a deletion at position 147. Furthermore, the 2009-like (blue) and 1991-like (orange) conformations showed periodic cross-reactivity to historical strains demonstrating the chronological reoccurrence of epitopes of limited variability (Fig 4B-G).

**Figure 4.**
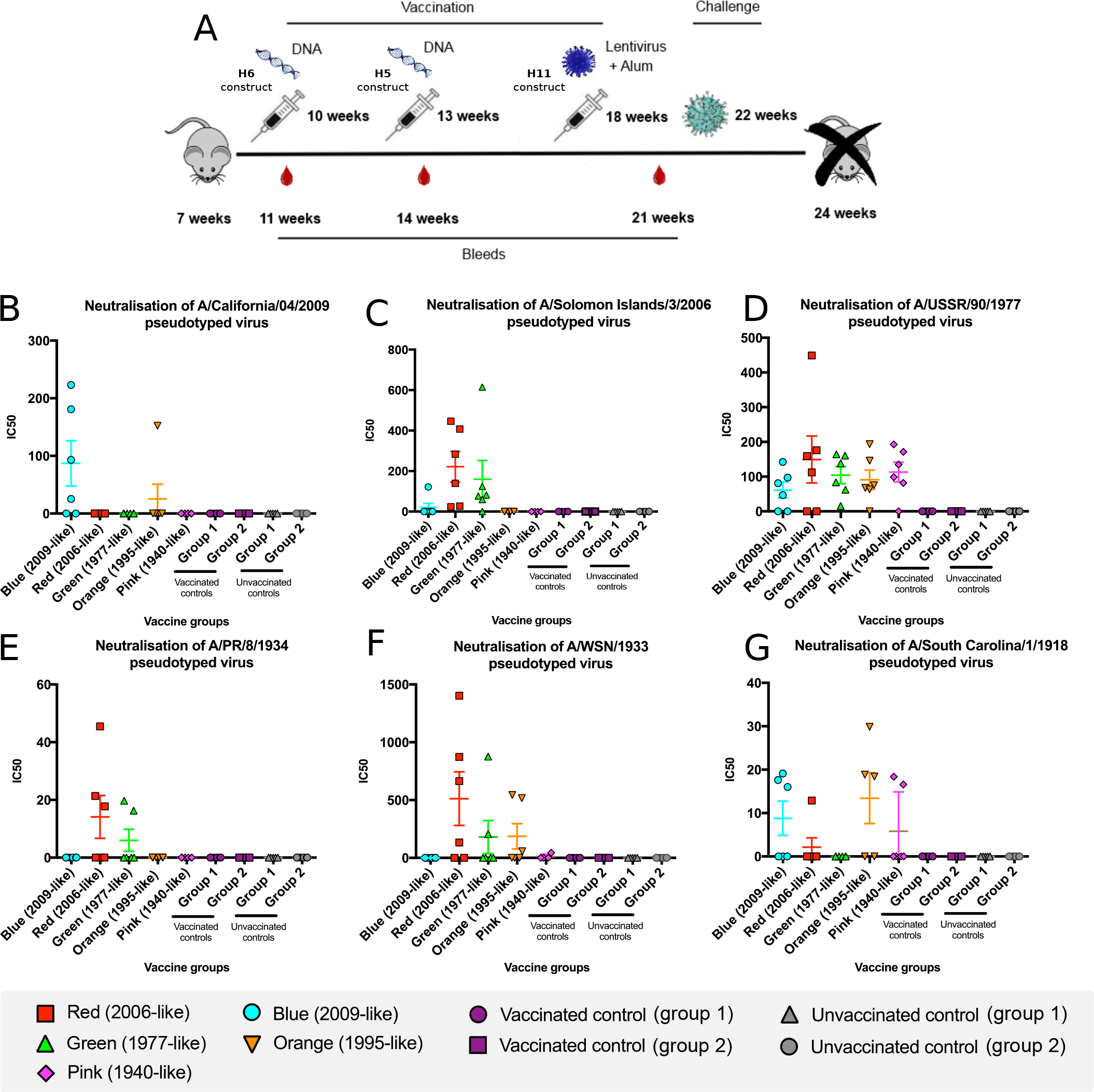
Sequential vaccination using chimeric HA constructs. (**A**) Five groups of mice were sequentially vaccinated with 2009-like (blue), 2006-like (red), 1995-like (orange), 1977-like (green) and 1940-like (pink) epitope sequences substituted into H6, H5 and H11 HAs (also see Table 1). Two further control groups were sequentially vaccinated with H6, H5 and H11 constructs without any sequence substituted into the HAs (vaccination controls). A further two groups were mock vaccinated (unvaccinated controls). (B-G) Pseudotype microneutralisation assays using 0.5 µl of sera from the bleed at 21 weeks. (H-I).

### Vaccination with epitopes from A/Solomon Islands/3/2006 and A/USSR/90/1977 protect against challenge with A/PR/8/1934

To test whether antibodies directed against these epitopes conferred protective immunity, the 2006-like (red) and 1977-like (green) epitope vaccinated groups were challenged with a strain collected in 1934 (A/PR/8/1934) (Fig 5A&C). The 2009-like (blue), 1995-like (orange) and 1940-like (pink) groups were challenged with a 2009 pandemic strain, A/California/04/2009 (Fig 5B&D). In each challenge experiment, an unvaccinated group (n=6) was included as well as a group vaccinated via the DNA-DNA-pseudotyped lentivirus regimen with the H6, H5 and H11 HAs without the substituted epitope conformations (n=6). Vaccination with the 2006-like (red) and 1977-like (green) epitope conformations conferred immunity to challenge with the A/PR/8/1934 virus (Fig 5A&C). As expected, vaccination with the 2009-like epitope also conferred immunity to challenge with A/California/04/2009 strain, which last circulated in 2009 (Fig 5B&D). These results demonstrate that epitopes that circulated in the A/Solomon Islands/8/2006 and A/USSR/90/1977 strains, which last circulated in 2006 and 1977 respectively, were able to produce antibodies that confer protection against challenge with the A/PR/8/1934 strain, which last circulated in 1934.

**Figure 5.**
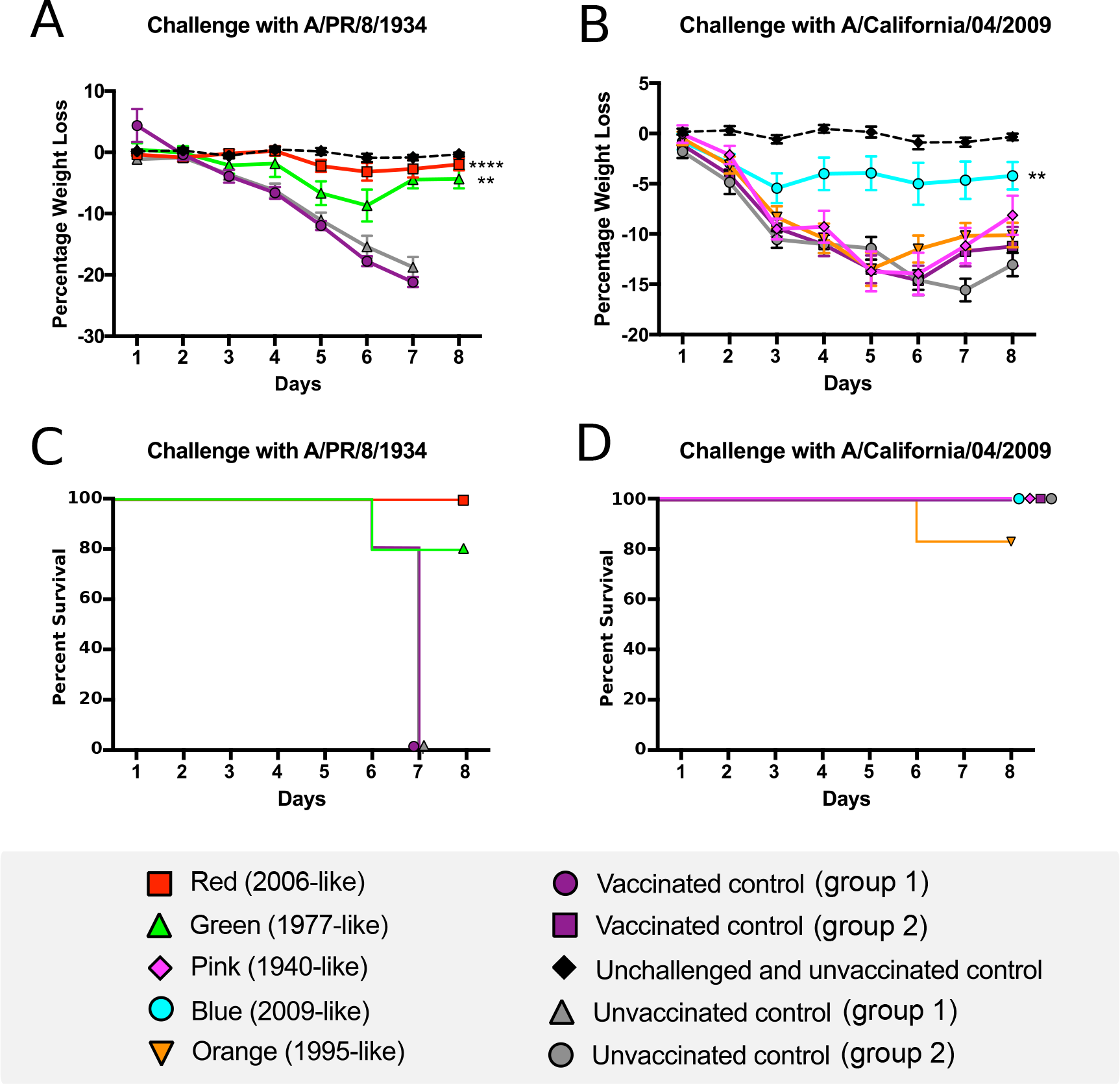
Influenza challenge of vaccinated mice with either A/PR/8/1934 or A/California/4/2009. (**A, B**) The graphs denote daily weight loss of the mice during the challenge. Mice of the same age, which were not vaccinated or challenged, are shown for reference and denoted ‘unchallenged and unvaccinated’. (**C, D**) Survival curve denoting the number of mice in each group. Mice were euthanised at 20% weight loss.

## Discussion

Our results demonstrate the existence of a highly immunogenic epitope of limited variability in the head domain of the H1 HA, which have been theorised by mathematical modelling studies to drive the antigenic evolution of influenza^9,10^. Sera from children aged 6-12 years taken in 2006/7 was shown to cross-react with a panel of historical isolates, the majority of which they will not have experienced (Fig 1A). This cross-reactivity was removed by mutagenesis of an epitope of limited variability identified through a structural bioinformatic analysis (Fig1B-F&2). We were further able to reproduce the cross-reactivity exhibited in the serology studies in a mouse model, and demonstrated that vaccination with the epitope conformations circulating in 2006 or 1977 induced protective immunity to challenge with a strain that last circulated in 1934 (Fig 4). Vaccination with other conformations of the epitope produced complementary but asynchronous cross-reactivity to historical strains (Figs 4&5). Furthermore, between 1918 and the present day, the 2006-like 147-deleted conformation of the epitope has occurred five times. In two instances, when circulating strains contained the 147-deleted conformation of the epitope, lineage replacement of the H1N1 strain occurred (in 1957 and 2008). This suggests that the possession of a conformation of the epitope in which 147 is present conferred a very significant selective advantage once population immunity has built up against the 147-deleted conformation.

Another site of limited variability identified by our analysis in the head domain of H1 HA appears to be centred on position 180 (linear numbering, position 166: H3 numbering, position 163: WHO numbering). Linderman et al, 2014 and Huang et al, 2015 have identified and purified antibodies that bind to a site including position 180 and neutralise A/California/04/2009, A/USSR/07/1977 and A/Brevig Mission/1/1918 but not A/Solomon Islands/30/2006, A/New Caledonia//1999 and A/PR/8/1934, displaying cyclical cross-reactivity in a similar but asynchronous manner to OREO^19,20,22^. We have identified similar patterns of cross-reactivity to historical isolates in young children aged 12-18 months whose sera was collected in 2013 (Fig S1B). However, as this site is periodically covered by glycosylation, the ‘OREO’ epitope is likely to be a better vaccine target.

Currently available influenza vaccines are believed to target epitopes of very high variability on the haemagglutinin and neuraminidase surface glycoproteins. This requires them to be continuously updated, with the only alternative being seen as the artificial boosting of immunity to conserved epitopes of low immunogenicity. By identifying such epitopes, we have established an alternate method of producing improved influenza vaccines: targeting highly immunogenic epitopes of limited variability as opposed to targeting highly immunogenic epitopes of high variability or conserved epitopes of low immunogenicity. Through vaccination against the various conformations of the epitope of limited variability identified in this study, it is possible to induce immunity to all previous and future H1N1 strains. Alternatively, directing immunity against a smaller number of the identified epitope conformations will reduce the frequency with which the seasonal influenza vaccine requires updating. The evolutionary framework on which these studies are based^9,34^ applies generally to other subtypes of influenza A such as H3N2 ^34^ and also to influenza B, suggesting that epitopes of limited variability can also be identified in these viruses. Consequently, the same strategy could be used to produce vaccines for other subtypes of human influenza, as well as swine and avian influenza viruses, and potentially other viruses.

## Materials and Methods

### Serum samples

88 serum samples from young children aged 6 to 12 years were collected in Oxford in 2006/2007. All donors gave written informed consent for research use of blood samples with ethical approval by a local research ethics committee 16/SC/0141.

### Enzyme-linked immunosorbant assay ELISA

Anti HA1 antibody responses were measured using ELISAs. In brief, Nunc-Immuno 96 well plates were coated with 1.0µg ml^-1^ of HA1 protein Sino Biological Ltd, China in PBS buffer and left overnight at 4°C. Plates were washed 6x with PBS-Tween PBS/T, then blocked with casein in PBS for 1 hour at room temperature RT. Serum or plasma was diluted in casein-PBS solution at dilutions ranging from 1:50 to 1:1000, before being added to Nunc-Immuno 96 well plates in triplicate. Plates were incubated at 4°C overnight before being washed as previously described. Secondary antibody rabbit anti-human whole IgG conjugated to alkaline phosphatase, Sigma was added at a dilution of 1:3000 in casein-PBS solution and incubated for 1.5 hours at RT. After a final wash, plates were developed by adding 4-nitrophenyl phosphate substrate in diethanolamine buffer Pierce, Loughborough, UK and optical density OD was read at 405nm using an ELx800 microplate reader Cole Parmer, London, UK. A reference standard comprising of pooled cross-reactive serum and naïve serum on each plate served as a positive and negative control respectively. Data is presented as arbitrary units determined using the NIH standard calculator based on subtraction of background from samples and interpolation from the standard curve using a 4 parameter fit model^35^.

A positive reference standard was used on each plate to produce a standard curve. The standard was made from cross-reactive serum against each HA1 protein. It was added in duplicate at an initial dilution of 1:100 in casein-PBS solution and diluted 2-fold 10 times, starting with an arbitrary value of antibody units determined using the NIH standard calculator. Three blank wells containing casein-PBS solution only, and a further 3 blank wells containing naïve human sera or plasma were used as negative controls. The mean of the OD values of the naïve sera was then subtracted from all OD values on each plate before triplicates were fitted to a 4 parameter standard curve using the positive reference standard.

### Pseudotyped influenza virus production

Pseudotyped lentiviruses displaying influenza HAs were produced by transfection of HEK 293T/17 cells with 1.0 µg of gag/pol construct, p8.91, 1.5 µg of a luciferase reporter carrying construct, pCSFLW, 250 µg of TMPRSS4 expressing construct and 1.0 µg of HA glycoprotein expressing construct Temperton et al 2007. Transfections were performed in 10 ml of media DMEM 10% FCS, 1% pencillicin-stephomycin, 20% L-gluatmate and left for 8 hours. 1 unit of endoengeous NA Sigma, US was added to 10 ml of new media to induce virus budding. Media was removed 48 hours post-induction of budding and filtered with a 0.45 µm syringe. The pseudotyped influenza viruses were stored at −80°C.

### Pseudotyped influenza virus titration

Serial dilutions were made of pseudotyped influenza virus preparations in Corning Costar plates 96-well plates Promega, USA.10^4^ HEK 293T/17 cells were added to each well and incubated for 3 days at 37°C. The cells were then lysed with BrightGlo reagent Promega, USA and the relative light units of the cell lysate determined using a Varioscan luminometer microplate reader Thermo Fisher Scientific, USA.

### Pseudotype microneutralisation assay

Neutralising antibodies were quantified using a pseudotype microneutralisation assay. Serially diluted sera was added to Corning Costar plates 96-well plates Promega, USA, before being incubated with 10^6^ RLU pseduotyped influenza virus for 1 hour at 37°C. Typically, 1 µl of sera was used per assay, however for comparison of WT and SDM pseudotyped influenza viruses, 5 µl of sera was used. HEK 293T/17 cells 2.0*10^5 cells ml^-1^ were subsequently added to each well and incubated for 3 days at 37°C. The cells were lysed with BrightGlo reagent Promega and the relative light units of the cell lysate determined using a Varioscan luminometer microplate reader Thermo Fisher Scientific, USA. The reduction of infectivity was determined by comparing the RLU in the presence and absence of antibodies and expressed as percentage neutralisation. The 50% inhibitory dose IC50 was defined as the sample concentration at which RLU were reduced 50% compared to virus control wells after subtraction of background RLU in cell only control wells. At least two replicates were performed for each biological sample to ensure replicability.

### Structural bioinformatic analysis

Amino acids present on the surface of various H1 HAs were determined by calculating the accessibility of amino acids on the surface of the crystal structures of A/Brevig Mission/1/1918 1RUZ, Gamblin et al, 2004, A/Puerto Rico/8/1934 1RU7, Skehel et al, 2004 and A/California/4/2009 3LZG, Xu et al, 2010 and A/Washington/5/2011 4LXV, Yang et al, 2014 HAs using Swiss-Pdb viewer. Areas of 600 Å^2^, 800 Å^2^ and 1000 Å^2^ were mapped onto the surface of the crystal structures by determining the distances between the α carbon of a given amino acid and all others within a structure. Those residues whose α carbon sequences were within the specified area were recorded and used to produce disrupted peptide sequences for a given binding site. Antibody binding site variability was calculated as the mean pairwise hamming distance between the consensus sequences collected between 1918 to 2016. The sequences were aligned using MUSCLE before being manually curated using AliView.

### Vaccination of mice

All animal work was approved by the University of Oxford Animal Care and Ethical Review Committee. Balb/c mice n=6 Envigo, UK were sequentially vaccinated with the OREO sequences substituted into H6, H5 and H11 HA backbones in a prime-boost-boost regime at intervals of 3-4 weeks. As a backbone control, two groups were vaccinated with native H6, H5 and H11 constructs. A further two groups were mock vaccinated and used as an unvaccinated control. The prime and first boost were administered as a 100 µg *intra muscular* injection of DNA into the *musculus tibialis*, whilst the final vaccination was administered as an *intra muscular* injection into the *musculus tibialis* of 8 HI units of lentivirus pseudotype displaying the chimeric H11 HA in Alum adjuvant Alhydrogel, Invitrogen, USA at a 1:1 volume ratio.

### Haemaggutinin inhibition assay

Pseudotyped lentivirus displaying influenza HA were diluted 2-fold down a 96 well plate and mixed with 50 µl of 4% chicken red blood cells. After an hour the coagulation of red blood cells was assessed visually to determine the point at which coagulation could no longer be observed.

### Influenza challenge

At 5 weeks post final boost mice were challenged *intra nasally* with either A/PR/8/1934 at 1.0 x 10^3^ PFU per mouse or A/California/04/2009 at 1.5 x 10^5^ PFU per mouse. Mice were weighed daily. At 3 days post challenge food in proportion to the number of mice in each cage was placed on the floor of the cage. Mice were euthanised at their pre-determined humane end point 20% weight loss or if they showed no sign of recovery at 7 days post-challenge.

### Statistical analysis

Student’s *t*-tests were performed to determine all p-values shown in Figure 1.

Area under the curve was calculated for the mouse weight loss data (Fig 5A&C, main text), and analysed in a single-factor ANOVA. Between-group comparisons were then performed using Tukey’s post hoc method for pairwise comparison correction to provide corrected p-values.

Fisher’s Exact test was used to determine survival differences in the experimental groups after seven days (Fig 5B&D). All p-values were adjusted to multiple comparisons using the Bonferroni-Holm correction.

### Phylogenetic analysis

RAxML version 8.2.11 was used to build a maximum likelihood tree based on the strain HA amino acid sequences, using a gamma distributed site heterogeneity model and the amino acid FLU substitution model. Tip-to-root distance was regressed against sequence dates, using a best fitting root, in Tempest V1.5.1. This yielded an R-squared of 0.886 and 0.834 for the ≤2008 and ≥2009 data, respectively, indicating a good fit between the genetic distance and the time of sampling. The colour of branches was determined by the identify of amino acids at positions 147 and 158, which are the variable amino acids at the centre of the amino acid binding site. Blue OREO was defined as 147 K, 158 no lysine; orange OREO as 147 lysine and 158 lysine; green OREO as 147 arginine; red OREO as 147 deletion; pink OREO 147 isoleucine.

Accession numbers can be found in the supplementary material.

## Data availability statement

The datasets generated during and/or analysed during this current study are available from the corresponding author on reasonable request.

## Code availability statement

The code will be made available to anybody on request

## Acknowledgements

We would like to thank Dr John S Tregoning Imperial College for kindly providing us with the viruses for the influenza challenge. We thank the parents/guardians who gave written informed consent for use of these blood samples for research by the Oxford Vaccine Centre Biobank, with ethical approval by a local research ethics committee 16/SC/0141. Other ethical approvals were provided by the OXTREC review board. Funding for the study was provided by a Royal Society Translation Award, a MRC confidence in concept grant and an ERC Advanced grant (DIVERSITY).

## Supplementary Information

**Figure S1.**
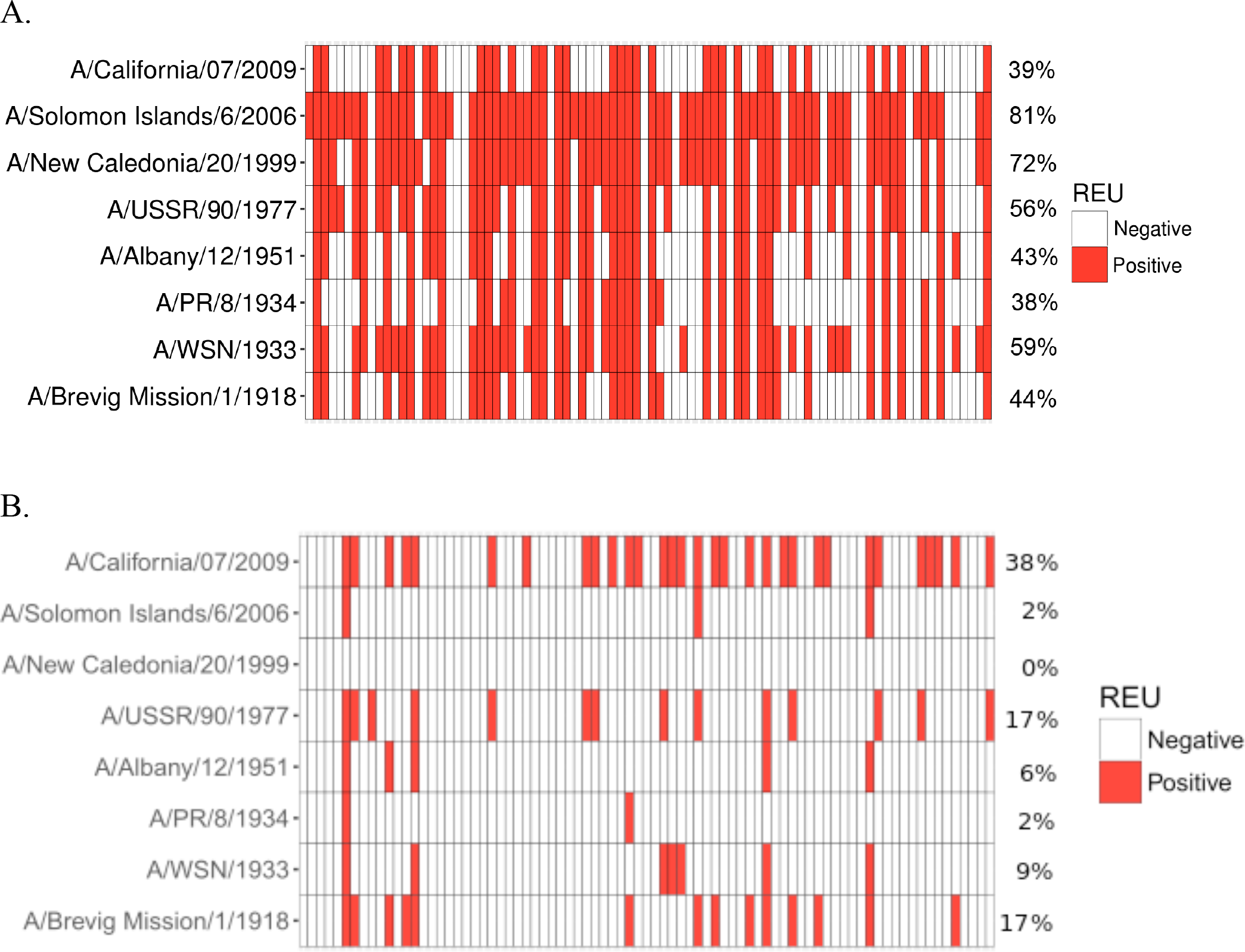
ELISA analysis of sera from children aged 6 to 12 years and 12 to 18 months using the HA1 domain of a number of chronologically dispersed strains. A. ELISA analysis of sera from children aged 6 to 12 years. B. ELISA analysis of sera from children aged 12 to 18 months. The HA1 domain of a number of chronologically dispersed strains was used as an antigen. All ELISAs were standardised using positive sera. Samples were either allocated as ‘positive’ (red) or ‘negative’ (white) based on whether they fell into the linear range of the standard curve. The percentage of the samples that displayed reactivity to each strain is provided on the right hand side of the figure.

**Figure S2.**
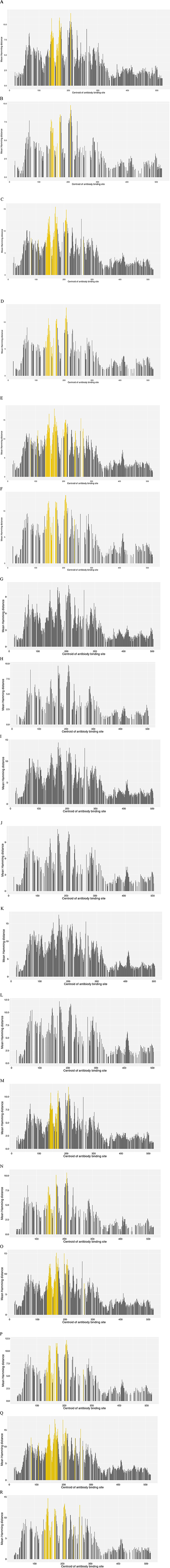
Variation in antibody binding sites mapped onto the surface of various H1 HA crystal structures. Variation in antibody binding sites mapped onto the surface of various H1 HA crystal structures. Variability in antibody binding sites ABS were mapped to the crystal structure of A/California/04/2009 to determine the variability of each antibody binding site based on parameters of (**A**) 600 Å^2^, >1% accessibility (**B**) 600 Å^2^, >10% accessibility, (**C**) 800 Å^2^, >1 accessibility (**D**) 800 Å^2^, >10% accessibility (**E**) 1000 Å^2^, >1% accessibility (**F**) 1000 Å^2^, >10% accessibility. Variability in antibody binding sites ABS were mapped to the crystal structure of A/PR/8/1934 to determine the variability of each antibody binding site based on parameters of (**G**) 600 Å^2^, >1% accessibility (**H**) 600 Å^2^, >10% accessibility, (**I**) 800 Å^2^, >1 accessibility (**J**) 800 Å^2^, >10% accessibility (**K**) 1000 Å^2^, >1% accessibility (**L**) 1000 Å^2^, >10% accessibility. Variability in antibody binding sites ABS were mapped to the crystal structure of A/Brevig Mission/1/1918 to determine the variability of each antibody binding site based on parameters of (**M**) 600 Å^2^, >1% accessibility (**N**) 600 Å^2^, >10% accessibility, (**O**) 800 Å^2^, >1 accessibility (**P**) 800 Å^2^, >10% accessibility (**Q**) 1000 Å^2^, >1% accessibility (**R**) 1000 Å^2^, >10% accessibility. Accessibility is determined by the accessibility of an amino acid to a water molecule. The amino acid position at the centre of each centroid is used to denote the ABS. In each instance ABS including position 147 are highlighted in yellow. Similar analyses were performed for the A/Washington/05/2011 crystal structure producing a similar pattern to the A/California/04/2009 analysis data not shown. ABS highlighted in yellow include position 147. Equivalents of these ABS can be found in A/PR/8/1934 but these have not been highlighted as there is a deletion in position 147.

**Figure S3.**
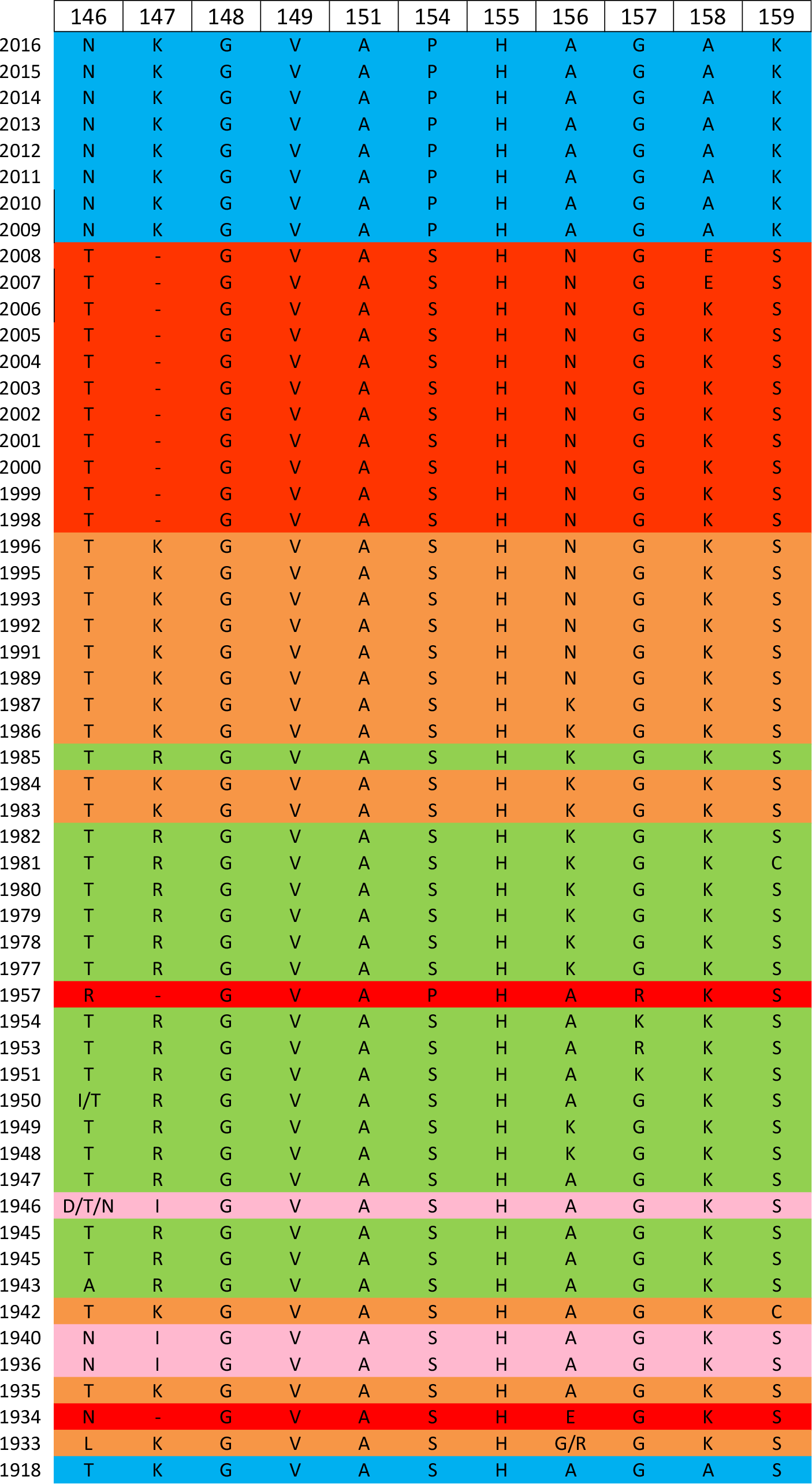
Identification of the various conformations of a site of limited variability in the head domain of the H1 HA through structural bioinformatic analysis. Analysis of consensus sequences corresponding to the disrupted peptide sequence of the antibody binding site of lowest variability containing position 147 OREO in the >10%, 1000 Å^2^ plot for A/California/4/2009.

**Table S1.**
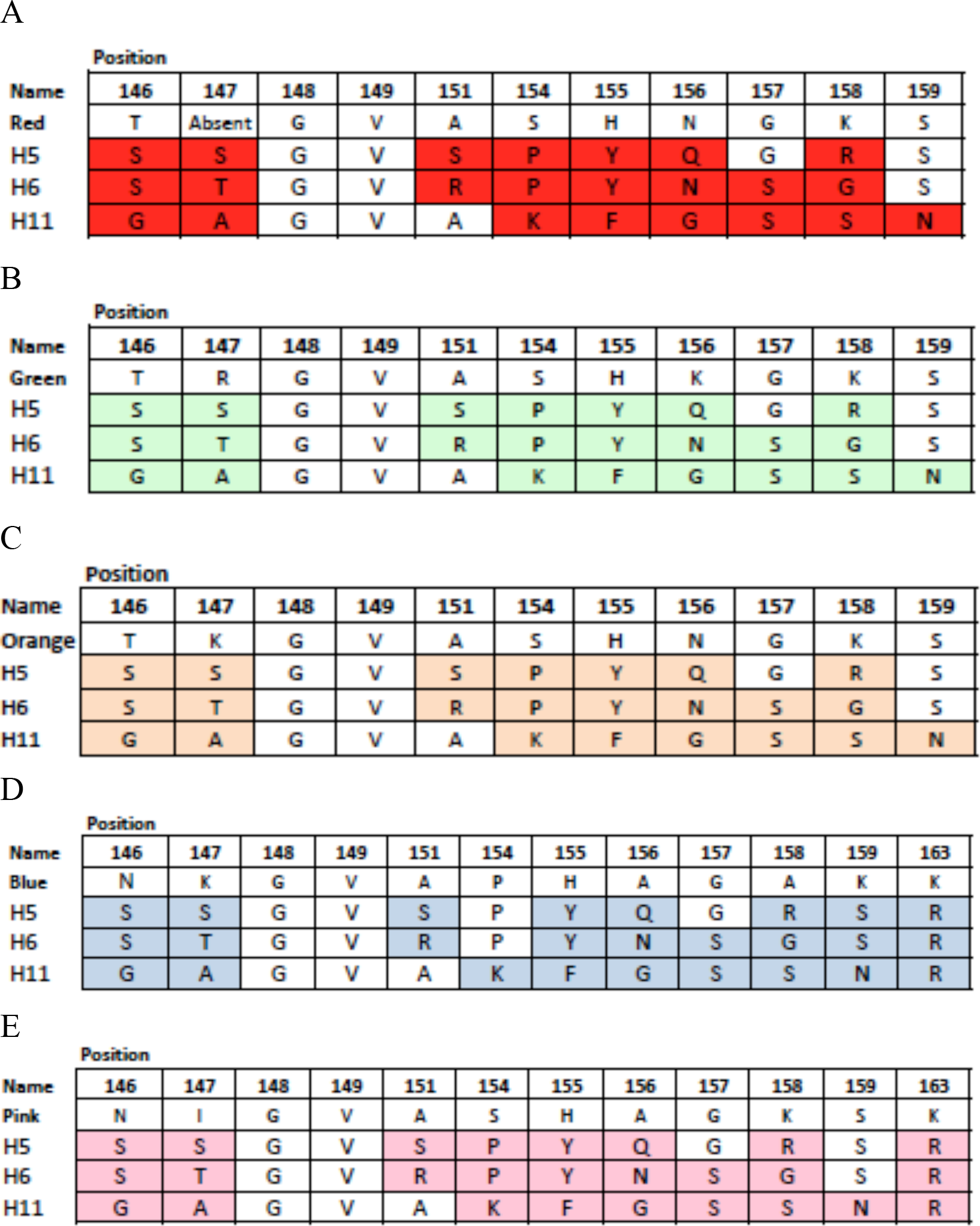
Sequences cloned into H6, H5 and H11 HAs. Amino acids changed in the H5, H6 and H11 backbones to the (**A**) red conformation 2006 H1 HA sequence, (**B**) green conformation (1977 sequence), (**C**) orange conformation (1995 sequence), (**D**) blue conformation (2009 sequence) and (**E**) pink conformation (1940 sequence).

|  | Position |  |  |  |  |  |  |  |  |  |  |
| --- | --- | --- | --- | --- | --- | --- | --- | --- | --- | --- | --- |
| Name | 146 | 147 | 148 | 149 | 151 | 154 | 155 | 156 | 157 | 158 | 159 |
| Red | T | Absent | G | V | A | S | H | N | G | K | S |
| H5 | S | S | G | V | S | P | Y | Q | G | R | S |
| H6 | S | T | G | V | R | P | Y | N | S | G | S |
| H11 | G | A | G | V | A | K | F | G | S | S | N |

B
|  | Position |  |  |  |  |  |  |  |  |  |  |
| --- | --- | --- | --- | --- | --- | --- | --- | --- | --- | --- | --- |
| Name | 146 | 147 | 148 | 149 | 151 | 154 | 155 | 156 | 157 | 158 | 159 |
| Green | T | R | G | V | A | S | H | K | G | K | S |
| H5 | S | S | G | V | S | P | Y | Q | G | R | S |
| H6 | S | T | G | V | R | P | Y | N | S | G | S |
| H11 | G | A | G | V | A | K | F | G | S | S | N |

C
|  | Position |  |  |  |  |  |  |  |  |  |  |
| --- | --- | --- | --- | --- | --- | --- | --- | --- | --- | --- | --- |
| Name | 146 | 147 | 148 | 149 | 151 | 154 | 155 | 156 | 157 | 158 | 159 |
| Orange | T | K | G | V | A | S | H | N | G | K | S |
| H5 | S | S | G | V | S | P | Y | Q | G | R | S |
| H6 | S | T | G | V | R | P | Y | N | S | G | S |
| H11 | G | A | G | V | A | K | F | G | S | S | N |

D
|  | Position |  |  |  |  |  |  |  |  |  |  |  |
| --- | --- | --- | --- | --- | --- | --- | --- | --- | --- | --- | --- | --- |
| Name | 146 | 147 | 148 | 149 | 151 | 154 | 155 | 156 | 157 | 158 | 159 | 163 |
| Blue | N | K | G | V | A | P | H | A | G | A | K | K |
| H5 | S | S | G | V | S | P | Y | Q | G | R | S | R |
| H6 | S | T | G | V | R | P | Y | N | S | G | S | R |
| H11 | G | A | G | V | A | K | F | G | S | S | N | R |

**Table S1: Sequences cloned into H6, H5 and H11 HAs.**
|  | Position |  |  |  |  |  |  |  |  |  |  |  |
| --- | --- | --- | --- | --- | --- | --- | --- | --- | --- | --- | --- | --- |
| Name | 146 | 147 | 148 | 149 | 151 | 154 | 155 | 156 | 157 | 158 | 159 | 163 |
| Pink | N | I | G | V | A | S | H | A | G | K | S | K |
| H5 | S | S | G | V | S | P | Y | Q | G | R | S | R |
| H6 | S | T | G | V | R | P | Y | N | S | G | S | R |
| H11 | G | A | G | V | A | K | F | G | S | S | N | R |

**Table S2.**
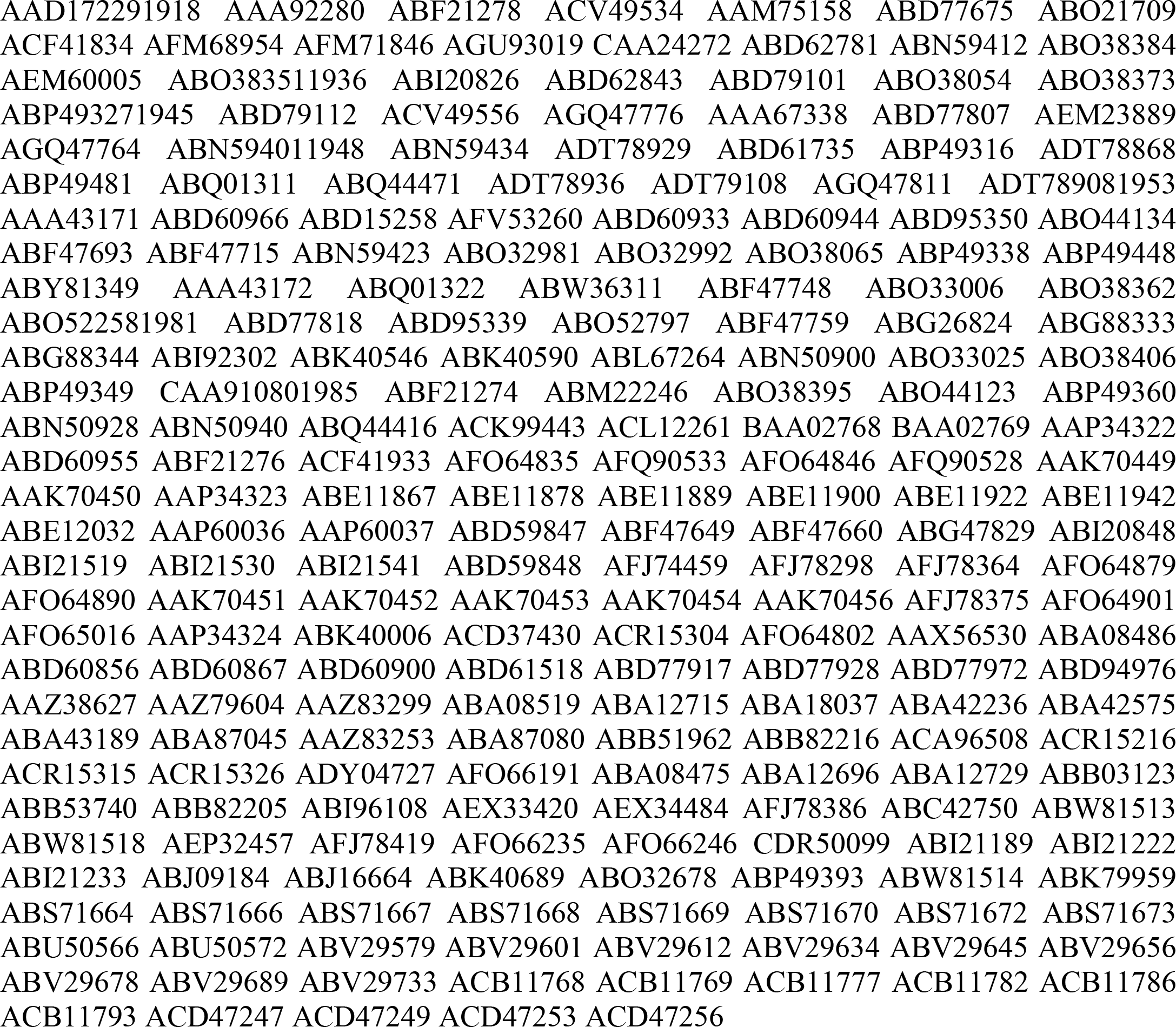
Accession numbers pre-2009 pandemic.

**Table S3.**
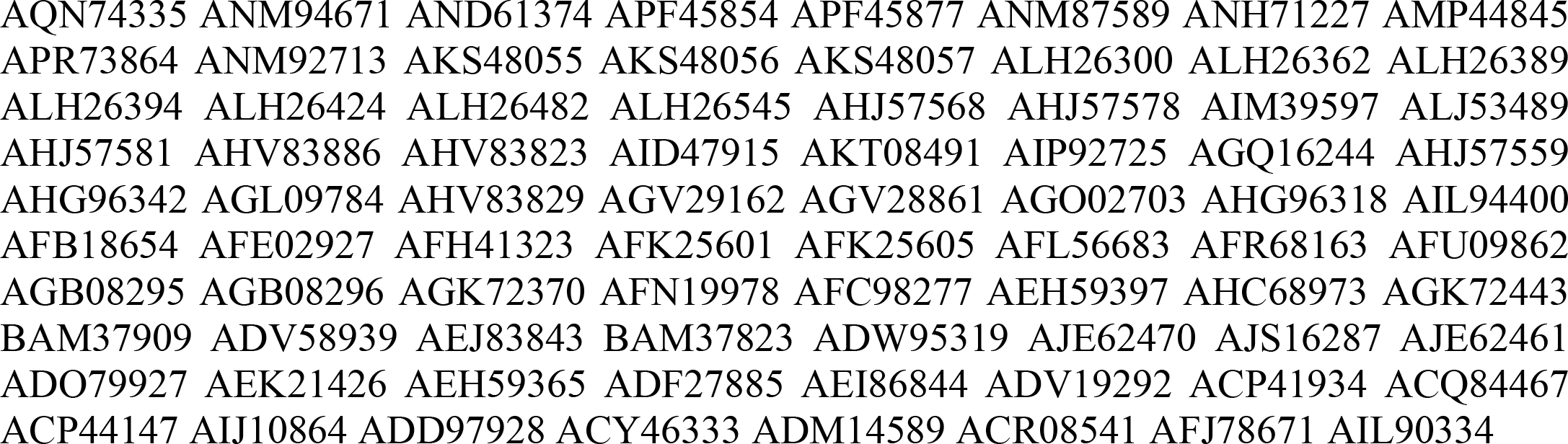
Accession numbers post-2009 pandemic.

